# An inductive, supervised approach for predicting gene–disease associations using phenotype ontologies

**DOI:** 10.1101/2025.05.07.652682

**Authors:** Safana Bakheet, Fernando Zhapa-Camacho, Robert Hoehndorf

## Abstract

**Motivation:** Predicting gene–disease associations (GDAs) is the problem to determine which gene is associated with a disease. The problem can be framed as a ranking problem where genes are ranked based on a set of phenotypes using a measure of phenotype similarity. When phenotypes are described using phenotype ontologies, ontology-based semantic similarity measures are used. Traditional semantic similarity measures use only the ontology taxonomy. Recent methods based on ontology embeddings compare phenotypes in latent space; these methods can use all ontology axioms as well as a supervised signal, but are inherently transductive, i.e., queries must already be known at the time of learning embeddings, and therefore these methods do not generalize to novel diseases (sets of phenotypes) at inference time.

**Results:** We developed an inductive method for ranking genes based on a set of phenotypes. Our method first uses a graph projection to map axioms from phenotype ontologies to a graph structure, and then uses ontology embeddings to create latent representations of phenotypes. We use an explicit aggregation strategy to combine phenotype embeddings into representations of genes or diseases, allowing us to generalize to novel sets of phenotypes. We also develop a method to make the phenotype embeddings and the similarity measure task-specific by including a supervised signal from known gene–disease associations. We apply our method to mouse models of human disease and demonstrate that we can significantly improve over inductive baseline measures, and reach a performance similar to transductive methods for predicting gene–disease associations while being more general.

**Availability and Implementation:** https://github.com/bio-ontology-research-group/ISGDA

## Introduction

Gene–disease associations (GDAs) for Mendelian diseases can be identified through multiple computational approaches. Mendelian diseases, characterized by single-gene mutations following Mendelian inheritance patterns, are predominantly rare disorders affecting fewer than 1 in 2,000 individuals (Umair and Waqas, 2023). Computational methods for identifying these associations include: (1) guilt-by-association methods that leverage biological networks where genes with similar network properties to known disease genes are implicated in similar diseases (Köhler *et al*., 2008) (2) phenotype similarity to databases of patients or diseases that connects patients with similar phenotypic profiles to known genetic causes (Köhler *et al*., 2009) and (3) phenotype similarity to model organisms, which provides substantially more data for inference (Hoehndorf *et al*., 2011). This last approach is particularly useful as wet lab validation of gene–disease associations remains time-consuming and expensive (Nunes, 2021), while computational methods can efficiently leverage large gene– phenotype and disease–phenotype datasets (Yuan *et al*., 2022). For rare Mendelian diseases, these approaches are crucial for advancing diagnosis and treatment options for affected patients. Phenotypes are recorded using standardized ontologies that enable computational analysis. Human phenotypes are described using the Human Phenotype Ontology (HPO) (Gargano *et al*., 2024), while mouse phenotypes are captured in the Mammalian Phenotype Ontology (MP) (Smith and Eppig, 2009). These species-specific ontologies structure phenotypes hierarchically with formal logical definitions. Cross-species ontologies such as UPheno (Matentzoglu *et al*., 2024) and PhenomeNET (Hoehndorf *et al*., 2011) facilitate comparisons between human and mouse phenotypes by relating classes of phenotypes in different species axiomatically and thereby making them comparable.

Phenotypes are then used to predict gene–disease associations through a ranked retrieval approach. This process involves using a phenotype similarity measure (a semantic similarity measure) to query a database of genotype–phenotype associations, e.g., from the Online Mendelian Inheritance in Men (OMIM) (Amberger *et al*., 2015) or the Mouse Genome Informatics (MGI) (Baldarelli *et al*., 2024) databases. Genotypes, usually representing loss of function of one or two alleles of a gene, are then ranked based on their phenotype similarity to the query disease. This ranking enables prioritization of candidate genes that are most likely to be causally related to a disease (Gkoutos *et al*., 2018). The effectiveness of this approach relies on the accuracy of the phenotype similarity measure and how complete the underlying phenotype data is.

Traditional semantic similarity measures for phenotype comparison are typically hand-crafted and can generalize to novel phenotype sets for querying. Examples include Resnik’s information content-based measure (Resnik, 1995) combined with the Best Match Average (BMA) approach for combining multiple comparisons (Harispe *et al*., 2015). Resnik quantifies similarity based on the information content of the most informative common ancestor of two classes in an ontology; the BMA then combines multiple (pairwise) similarities into a single similarity score. It is this aggregation strategy that makes similarity measures like Resnik able to generalize to novel, unseen diseases. Other similarity measures and aggregation strategies include the weighted Jaccard index (Pesquita *et al*., 2008) which also generalizes to novel combinations of classes.

These measures have been successfully applied to the gene–disease association prediction tasks (Putman *et al*., 2024; Alghamdi *et al*., 2022; Hoehndorf *et al*., 2011; Smedley *et al*., 2013; Chen *et al*., 2012). However, semantic similarity measures primarily rely on the phenotype ontology’s hierarchical structure and do not consider other axioms between phenotypes. Furthermore, because semantic similarity measures are hand-crafted, they do not adapt to the data or task of predicting gene–disease associations. More recent work has applied machine learning to generate embeddings of phenotypes, genes, and diseases. These knowledge graph or ontology embeddings (Chen *et al*., 2025) learn latent representations of single phenotypes or sets of phenotypes, which can then be used either through a vector similarity measure (e.g., cosine similarity) to perform ranked retrieval, or using a supervised method like a learning-to-rank approach with a neural network (Chen *et al*., 2021b). Embedding-based methods applied to the task of predicting gene–disease associations based on phenotype similarity include Onto2Vec (Smaili *et al*., 2018) and OPA2Vec (Smaili *et al*., 2019), DL2Vec (Chen *et al*., 2021a), OWL2Vec* (Chen *et al*., 2021a), and SmuDGE (Alshahrani and Hoehndorf, 2018).

Embedding-based approaches are inherently transductive. Transductive learning requires that all entities (diseases, genes) that will be used during inference must already be available during the training phase. This means that these models cannot generalize to previously unseen diseases without complete retraining, thereby limiting their applicability to patients with a rare but previously uncharacterized disease. Furthermore, the embedding methods that were applied to the task of predicting gene–disease associations only improve predictive performance over traditional semantic similarity measures like Resnik’s measure when they incorporate a supervised signal — known gene–disease associations — during training. This supervised signal may introduce bias as the model can “memorize” known associations rather than learning generalizable patterns from phenotype data alone (Alghamdi *et al*., 2022), or predict entirely based on the number of times a certain disease or gene was seen during training (Smaili *et al*., 2019).

The limitations of transductive approaches extend beyond basic gene–disease association prediction to variant prioritization applications. Systems such as Exomiser (Robinson *et al*., 2014) or EmbedPVP (Althagafi *et al*., 2024) combine phenotype-based gene–disease association prediction with variant pathogenicity measures to prioritize potentially causal variants in clinical settings. For such applications, the ability to make inductive predictions — generalizing to novel patients with previously unseen combinations of phenotypes — is even more critical. Patients with rare or previously uncharacterized genetic conditions cannot benefit from approaches that require prior knowledge of their specific disease during model training. While systems like Exomiser use semantic similarity, EmbedPVP, while showing a higher predictive performance, uses a transductive method for predicting gene– disease associations.

We developed a fully inductive method for gene–disease association prediction based on ontology embeddings while retaining a supervised learning component. Our approach enables ranking genes based on phenotype similarity without requiring the test diseases (sets of phenotypes) to be present during training. We find that our method outperforms traditional semantic similarity measures while maintaining the ability to generalize to previously unseen diseases.

Our main contributions include: (1) a novel inductive framework for gene–disease association prediction that generalizes to new combinations of phenotypes; and (2) an empirical validation of our method’s effectiveness compared to established semantic similarity measures.

## Materials and Methods

### Datasets

We obtained gene–phenotype associations from the MGI database (Baldarelli *et al*., 2024). Specifically, we used the file `MGI_GenePheno.rpt`, downloaded from the Mouse Genome Informatics Database on 10, July 2024. From this file we extracted gene identifiers and their corresponding phenotypes, encoded with the MP.

For disease–phenotype associations, we used the file `phenotype.hpoa` from the Human Phenotype Ontology database (Gargano *et al*., 2024), downloaded on 5 September 2024.

We used the UPheno cross-species phenotype ontology (Matentzoglu *et al*., 2024), downloaded on 21 August 2024. For all phenotype associations, we ensured that the phenotypes exist in the UPheno ontology, otherwise we omit the phenotype association.

To evaluate our ability to identify gene–disease associations, we used the file `MGI_Geno_DiseaseDO.rpt` from the Mouse Genome Informatics Database (Baldarelli *et al*., 2024), downloaded on 13 July 2024.

### Ontology preprocessing and graph projection

We define an ontology as the tuple 𝒪 = (Σ, *Ax*), where Σ = (**C, R, I**) provides the signature of the ontology (**C** is a set of classes, **R** is a set of roles and **I** is a set of individuals) and *Ax* provides the set of axioms over Σ. For example, axioms in *Ax* can follow the grammar *A* | *A* ⊓ *B* | *A* ⊔ *B* | ∃*r.A* | ∀*r.A* | *A*(*a*) | *r*(*a, b*)(Baader, 2003). We use the axioms in the UPheno ontology and added new axioms representing gene–phenotype, disease–phenotype and gene–disease associations. For example, for an association between a gene *g*_*i*_ and phenotype *p*_*j*_, we created the axiom *g*_*i*_ ⊑ ∃has phenotype.*p*_*j*_, which we added to UPheno. Similarly, disease–phenotypes associations were transformed to axioms *d*_*i*_ ⊑ ∃has symptom.*p*_*j*_ and disease–gene associations were added to UPheno as axioms *d*_*i*_ ⊑ ∃associated with.*g*_*j*_. A graph projection maps 𝒪 into a graph 𝒢 following a specific set of rules (Zhapa-Camacho and Hoehndorf, 2023). We projected UPheno and its extensions into graphs following the projection rules designed by OWL2Vec* (Chen *et al*., 2021a).

To evaluate the impact of different information sources on prediction performance, we constructed four distinct datasets with increasing amounts of information. Graph 1 contains only the UPheno ontology, with its phenotype classes for humans and mice. This baseline graph includes the hierarchical relationships between phenotype terms and the mappings between human and mouse phenotypes, but no gene or disease annotations. Graph 2 extends Graph 1 by adding gene– phenotype associations, connecting MGI gene identifiers to their associated MP terms. This graph incorporates information on which phenotypes are observed when specific mouse genes are mutated, but does not include disease information. Graph 3 further extends Graph 2 by adding disease–phenotype associations, connecting OMIM disease identifiers to their associated HPO terms. This graph contains the complete phenotypic profile of both genes and diseases, but does not include known gene–disease associations. Graph 4 contains a supervised signal and extends Graph 3 by adding known gene– disease associations between MGI genes and OMIM diseases (i.e., mouse models of human disease. This graph contains the complete information set, including the ground truth associations that serve as a supervision signal during training.

These four graph structures allow us to systematically evaluate how additional information affects the model’s ability to predict gene-disease associations. By comparing performance across these structures, we can determine whether the inclusion of gene–phenotype, disease–phenotype, or direct gene–disease associations has the greatest impact on prediction accuracy.

To ensure our method is truly inductive and can generalize to previously unseen diseases, we implemented a 10-fold cross-validation strategy based on disease splits. For Graphs 3 and 4, which contain disease entities, the disease set was randomly partitioned into 10 equally sized subsets. In each fold, 90% of diseases were used for training and 10% were held out for testing. This procedure guarantees that the diseases used for evaluation were never seen during the training phase, thus validating the model’s ability to make predictions on novel diseases based solely on their phenotypic profiles.

### Graph Embedding Models

We use several knowledge graph embedding methods (Wang *et al*., 2017) for our experiments, specifically TransE (Bordes *et al*., 2013), TransD (Ji *et al*., 2015), RotatE (Sun *et al*., 2019), and PairRE (Chao *et al*., 2021). Each embedding method captures different relational patterns in the knowledge graph. TransE uses the scoring function *f* (*h, r, t*) = − ∥*h* + *r* − *t*∥ and models relationships as translations in the embedding space. TransD employs *f* (*h, r, t*) = − *h*^⊥^ + *r* − *t*^⊥^, where entities are projected into relation-specific spaces, allowing more expressiveness for modeling complex relations. RotatE, with scoring function *f* (*h, r, t*) = − ∥*h* ° *r* − *t*∥, represents relations as rotations in complex vector space, enabling it to model various relation patterns including symmetry, antisymmetry, inversion, and composition. PairRE uses *f* (*h, r, t*) = − ∥*h* ° *r*_*H*_ − *t* ° *r*_*T*_ ∥ and employs dual relation-specific vectors to enhance expressiveness while maintaining computational efficiency.

### Semantic Similarity

Semantic similarity measures quantify the likeness of concepts based on their meaning and relationships within an ontology. For phenotype-based gene–disease association prediction, we use two approaches: Resnik’s information content-based measure and embeddings with cosine similarity.

Resnik’s semantic similarity measure (Resnik, 1995) quantifies the similarity between two ontology terms based on their shared information content (IC). For two phenotype terms p_1_ and p_2_, the similarity is defined as:

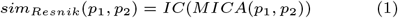

where *MICA*(*p*_1_, *p*_2_) is the most informative common ancestor of *p*_1_ and p_2_ in the ontology hierarchy, and *IC*(*p*) is the information content of term *p*, calculated as:

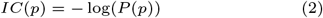

with *P* (*p*) representing the frequency of phenotype term *p* in the corpus of annotations. We used the Semantic Measures Library (Harispe *et al*., 2013, 2015) to compute Resnik semantic similarity.

For embedding-based similarity, we leverage the vector representations generated by knowledge graph embedding models. Given two phenotype terms with embeddings 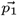 and 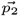, we calculate their similarity using cosine similarity:

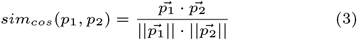

To compare sets of phenotypes associated with genes and diseases, we employ the Best-Match Average (BMA) approach. For a gene *g* with phenotypes *P*_*g*_ = {*p*_*g*1_, *p*_*g*2_, …, *p*_*gn*_} and a disease *d* with phenotypes *P*_*d*_ = {*p*_*d*1_, *p*_*d*2_, …, *p*_*dm*_}, the BMA score is:

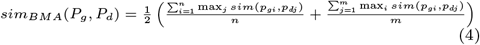

where sim can be either Resnik similarity or cosine similarity of embeddings. This approach finds the best possible match for each phenotype term in both sets and averages the results.

We note that phenotype terms are defined by the ontology and do not change. Therefore, the BMA method of computing similarity of sets of phenotypes is inherently inductive, i.e., can be applied to any (novel) set of phenotypes.

## Results

### Transductive prediction of gene–disease associations

We integrated a cross-species phenotype ontology, gene– phenotype associations resulting from single gene loss of function mutations in the mouse, and disease–phenotype associations from the Online Mendelian Inheritance in Men (OMIM) (Amberger *et al*., 2015), in a knowledge graph. We create two versions of this graph, the first one consisting of the phenotype ontology, gene–phenotype, and disease– phenotype associations. We call this graph “non-supervised”. We also included known gene–disease associations in the second version of this graph, which we call “supervised”. We then implemented a transductive approach that generates knowledge graph embeddings for the entities in the graphs, and uses the non-supervised and supervised graphs to directly predict gene–disease associations based on the similarity between embeddings of genes and diseases.

This transductive approach allows us to evaluate the performance of different embedding methods for GDA prediction and establish a comparison baseline for our inductive method and Resnik’s semantic similarity measure. In the transductive setting, gene–disease associations are predicted differently depending on the graph structure. For the non-supervised approach, which lacks direct gene–disease associations, predictions are made by computing cosine similarity directly between the embeddings of genes and diseases obtained from the knowledge graph. For the supervised graph, the model can leverage the gene–disease links present during training to make predictions during testing. In both cases, all genes and diseases used during testing must be present in the knowledge graph during training, making both approaches transductive by definition. We select 10% of diseases (with all their associated genes) for testing; for the supervised graph, we removed all gene–disease associations for these 10% diseases from the graph to ensure that the gene–disease associations on which we test are not present during training.

We evaluated four different knowledge graph embedding methods (TransE, TransD, RotatE, and PairRE) using this transductive approach. As shown in Table 1, TransD consistently outperformed the other embedding methods across all evaluation metrics. The large performance gap between TransD and the other embedding methods indicates that TransD’s entity projection mechanism is suitable for capturing the relationships in the knowledge graph we created. Based on these results, we selected TransD as the embedding method for all subsequent experiments.

**Table 1.** Performance comparison of different gene–disease association prediction methods. Results are reported for a 10% test set of diseases (one fold). Direct GDA scoring represents the transductive approach where genes and diseases must be present during training. BMA-based scoring represents the inductive approach that can generalize to new phenotype combinations. **Bolded** values are for the model that improves over the Resnik + BMA baseline and is the best-performing amongst other models. Note that for MR, lower is better.

| Method | Unsupervised |  |  |  |  | Supervised |  |  |  |  |
| --- | --- | --- | --- | --- | --- | --- | --- | --- | --- | --- |
|  | AUC | H@1 | H@10 | MR | MRR | AUC | H@1 | H@10 | MR | MRR |
| *Baseline* |  |  |  |  |  |  |  |  |  |  |
| Resnik + BMA | Same as Graph 4 |  |  |  |  | 0.9028 | 0.0635 | 0.2937 | 191.76 | 0.1461 |
| *Direct GDA Scoring (Transductive)* |  |  |  |  |  |  |  |  |  |  |
| TransE | 0.5200 | 0.0000 | 0.0159 | 696.87 | 0.0069 | 0.6099 | 0.0714 | 0.1508 | 236.48 | 0.1051 |
| TransD | 0.5856 | 0.0000 | 0.0159 | 384.23 | 0.0113 | **0.9156** | **0.3254** | **0.6190** | **62.48** | **0.4129** |
| RotatE | 0.3328 | 0.0000 | 0.0159 | 676.79 | 0.0082 | 0.8461 | 0.2063 | 0.3730 | 126.85 | 0.2712 |
| PairRE | 0.4237 | 0.0000 | 0.0159 | 663.05 | 0.0061 | 0.7291 | 0.0397 | 0.1508 | 250.79 | 0.0825 |
| *BMA-based GDA Scoring (Inductive)* |  |  |  |  |  |  |  |  |  |  |
| TransE | 0.6968 | 0.0000 | 0.0159 | 574.02 | 0.0097 | 0.7204 | 0.0159 | 0.0476 | 527.67 | 0.0318 |
| TransD | 0.8687 | 0.0397 | 0.2381 | 260.05 | 0.1025 | **0.9140** | **0.0556** | **0.3492** | **164.02** | **0.1447** |
| RotatE | 0.7118 | 0.0079 | 0.0079 | 523.79 | 0.0139 | 0.6820 | 0.0000 | 0.0556 | 482.67 | 0.0250 |
| PairRE | 0.7049 | 0.0000 | 0.0159 | 564.65 | 0.0131 | 0.7050 | 0.0000 | 0.0397 | 556.55 | 0.0126 |

When comparing the non-supervised and supervised graph, we also observe differences in prediction performance. The supervised approach shows overall higher performance across all metrics, indicating that the inclusion of gene–disease associations during training provides a useful signal for predicting new associations between genes and diseases already in the knowledge graph.

### Inductive Approach for Gene–Disease Association Prediction

In clinical settings, the gene–disease association prediction task frequently involves novel diseases or patients with previously uncharacterized conditions. While the set of genes remains stable, new diseases and phenotypic manifestations continue to be discovered, particularly for rare Mendelian disorders. This presents a fundamental limitation for transductive embedding approaches, which require diseases to be present during the training phase to generate their embeddings.

The key insight of our inductive approach is that while diseases may be novel, the phenotypes used to describe them are drawn from a stable, predefined ontology. Resnik’s semantic similarity measure is inherently inductive because it operates on phenotypes rather than directly on diseases or genes. It computes the best-match average (BMA) between sets of phenotypes associated with genes and diseases, making it applicable to any disease characterized by known phenotypes, even if the disease itself was not seen during training.

We extend this intuition to embedding-based approaches by computing BMA scores using cosine similarities between phenotype embeddings rather than directly comparing gene and disease embeddings. Given a gene *g* with phenotypes *P*_*g*_ and a disease *d* with phenotypes *P*_*d*_, we calculate the BMA score based on pairwise cosine similarities between phenotype embeddings. This approach allows us to predict associations for any disease described by phenotypes in our ontology, even if the disease itself was not included during training.

To evaluate our inductive approach, we conducted experiments using the TransD embedding model with four different graph structures of increasing complexity:

- Graph 1 includes only the UPheno ontology with cross-species phenotype mappings, providing a baseline that captures hierarchical and cross-species phenotype relationships but contains no gene or disease information.
- Graph 2 extends Graph 1 by adding gene–phenotype associations, enabling the model to learn relationships between genes and phenotypes while still lacking disease information. Note that the set of genes is always known and does not change, making this approach also inductive.
- Graph 3 further extends Graph 2 by incorporating disease– phenotype associations from OMIM. This graph contains the complete phenotypic profiles of both genes and diseases but does not include known gene–disease associations. As long as the disease (or set of phenotypes) which is tested is not included in this graph, predicting gene– disease associations with Graph 3 can also be inductive. We generate 10 versions of this graph, in each of which we remove 10% of all diseases; for inductive inference, we use the graph in which the test disease is not present.
- Graph 4, our supervised graph, extends Graph 3 by adding known gene–disease associations. Similarly to Graph 3, we implemented 10-fold cross-validation on diseases to ensure that test diseases were never seen during training.

Table 2 presents the performance of our inductive BMA-based approach across these graph structures. Several key findings emerge from these results: First, performance improves progressively with the addition of more information to the graph, with Graph 4 consistently outperforming the others. The supervised signal from known gene–disease associations significantly enhances the model’s ability to predict new associations. Second, for Graph 4, our inductive approach achieves an AUC of 0.8995, which is comparable to the transductive approach despite the more challenging task of generalizing to unseen diseases, i.e., inductive inference. This demonstrates that learning from phenotype patterns is nearly as effective as directly learning from gene–disease associations. Third, our inductive approach using TransD embeddings (AUC 0.8995) significantly improves over Resnik’s semantic similarity measure (p=0.0113, Mann-Whitney U test), confirming that learned phenotype embeddings can capture more complex relationships than handcrafted similarity measures based solely on information content, while still retaining the ability for inductive inference. These results validate that our approach remains effective when applied to diseases not seen during training, making it suitable for real-world clinical applications where newly characterized diseases (characterized as sets of phenotypes) are considered.

**Table 2.** Performance of different graph structures using TransD with the BMA-based approach. All results are based on 10-fold cross-validation for Graph 3 and Graph 4, ensuring test diseases were not seen during training. **Bold** values indicate best performance for each metric (note that for MR, lower is better).

| Methods with 10-fold, CV | AUC | H@1 | H@10 | MR | MRR |
| --- | --- | --- | --- | --- | --- |
| Resnik + BMA (baseline) | 0.8611 | 0.0564 | 0.2730 | 197.67 | 0.1248 |
| Graph 1 (Upheno only) | 0.7904 | 0.0655 | 0.2470 | 263.04 | 0.1259 |
| Graph 2 (Upheno + genes) | 0.7934 | 0.0624 | 0.2289 | 263.49 | 0.1156 |
| Graph 3 (Upheno + genes + diseases) | 0.8458 | 0.0564 | 0.2349 | 222.19 | 0.1142 |
| Graph 4 (Upheno + genes + diseases + GDAs) | **0.8995** | **0.0706** | **0.2849** | **154.55** | **0.1422** |

## Discussion

### Inductive Gene–Disease Association Prediction

Inductive approaches for gene–disease association prediction are critical for addressing the challenges of rare genetic disease diagnosis. The majority of Mendelian diseases are rare, with new conditions continually being characterized. Traditional transductive embedding approaches cannot handle previously unseen diseases without complete retraining, severely limiting their clinical utility. Our inductive method addresses this limitation by enabling predictions for novel diseases based solely on their phenotypic profiles.

This capability is particularly important when integrating gene–disease association prediction into variant prioritization systems such as Exomiser (Robinson *et al*., 2014) and PVP (Althagafi *et al*., 2024). These systems combine phenotype-based gene prioritization with variant pathogenicity metrics to identify causative variants in patients with genetic disorders. By adopting our inductive approach, these systems can handle patients with previously uncharacterized diseases or novel combinations of phenotypes without requiring prior knowledge of specific disease entities during model training.

Our approach extends the state-of-the-art in multiple ways. First, unlike traditional semantic similarity measures like Resnik’s, which rely on hand-crafted metrics, our method leverages knowledge graph embeddings to learn relationships between phenotypes across species. Second, unlike previous embedding approaches that require diseases to be present during training, our method operates at the phenotype level, making it inherently inductive. Third, we retain the benefits of supervised learning by incorporating gene–disease associations during training while maintaining the ability to generalize to unseen diseases. This balance between supervision and induction is likely the most significant advancement over existing approaches, and may open up possibilities for future improvements in predicting gene–disease associations.

### Embedding model performance

Our experimental results demonstrate that TransD consistently outperforms other embedding models (TransE, RotatE, and PairRE) for gene–disease association prediction. TransD’s entity projection mechanism, which maps entities into relation-specific spaces, appears particularly well-suited for capturing the relationships in our graph. This finding aligns with a previous study that also showed that models with relation-specific projections better handle heterogeneous knowledge graphs for gene–disease association prediction (Althagafi *et al*., 2024). More theoretical work is required in the future to understand exactly how TransD enables improved performance over other knowledge graph embedding methods.

The progressive improvement in performance with increasingly rich graph structures confirms the value of incorporating multiple information sources. While the UPheno ontology alone (Graph 1) provides a foundation for phenotype comparison, the addition of gene–phenotype associations (Graph 2), disease– phenotype associations (Graph 3), and known gene–disease associations (Graph 4) each contribute to improved predictive performance. This gradual enhancement demonstrates that each layer of biological knowledge adds signal that the embedding model can leverage. We also note that Graph 2 contains the information that can be used by the Resnik similarity measure, as information content is computed over genes; graphs 3 and 4 are able to utilize more information than available to Resnik’s measure.

Notably, the supervised model (Graph 4) achieves substantially higher performance than unsupervised alternatives, highlighting the importance of known gene–disease associations as a supervision signal. The need for a supervised signal has also been shown in all prior studies that rely on embeddings for computing gene–disease associations. However, unlike previous supervised approaches that sacrifice inductive capabilities, our method maintains the ability to generalize to unseen diseases through the BMA-based phenotype comparison.

### Comparison with Semantic Similarity Measures

Our comparison with Resnik’s semantic similarity measure reveals important insights about the strengths of embedding-based approaches. While Resnik’s measure has been widely used due to its simplicity and interpretability, it primarily relies on the information content of the most informative common ancestor in the ontology hierarchy. This approach neglects more complex relationships and cannot adapt to the specific patterns in gene–disease association data.

In contrast, our embedding-based approach learns latent representations that capture both hierarchical and non-hierarchical relationships in the ontology. The TransD model can identify axiom patterns in how phenotypes relate to each other across species, which may explain its superior performance. Additionally, by incorporating a supervised signal during training, our model can learn which phenotype patterns are most relevant for predicting gene–disease associations, rather than relying solely on general semantic similarity.

## Conclusions

Our study demonstrates that inductive, supervised gene– disease association prediction can successfully address the limitations of traditional approaches. By operating at the phenotype level rather than directly at the disease level, our method can generalize to novel diseases based solely on their phenotypic manifestations. This capability is particularly valuable for rare disease diagnosis where previously uncharacterized conditions regularly emerge.

The framework we have developed has implications for clinical genomics. By enabling accurate prediction of gene– disease associations for novel diseases, our approach can improve the prioritization of candidate genes in diagnostic settings. This can potentially reduce the time and cost of rare disease diagnosis, ultimately leading to earlier and more effective treatment interventions.

## Author contributions statement

S.B. designed and conducted experiments, wrote the software, analyzed and interpreted the results, and wrote the paper. F.Z.C. contributed to experiments and writing of code, and writing the paper. R.H. conceived of and supervised the work, acquired funding, and contributed to writing of the manuscript. All authors have read and critically revised the manuscript.

## Acknowledgements

This work has been supported by funding from King Abdullah University of Science and Technology (KAUST) Office of Sponsored Research (OSR) under Award No. URF/1/4675-01-01, URF/1/4697-01-01, URF/1/5041-01-01, REI/1/5235-01-01, and REI/1/5334-01-01. This work was supported by funding from King Abdullah University of Science and Technology (KAUST) – KAUST Center of Excellence for Smart Health (KCSH), under award number 5932, and by funding from King Abdullah University of Science and Technology (KAUST) – Center of Excellence for Generative AI, under award number 5940.

